# Exploring Energy Landscapes of Intrinsically Disordered Proteins: Insights into Functional Mechanisms

**DOI:** 10.1101/2021.01.28.428595

**Authors:** Antonio B. Oliveira, Xingcheng Lin, Prakash Kulkarni, José N. Onuchic, Susmita Roy, Vitor B.P. Leite

## Abstract

Intrinsically disordered proteins (IDPs) lack a rigid 3D structure and populate a polymorphic ensemble of conformations. Because of the lack of a reference conformation, their energy landscape representation in terms of reaction coordinates presents a daunting challenge. Here, our newly developed Energy Landscape Visualization Method (ELViM), a reaction coordinate-free approach, shows its prime application to explore frustrated energy landscapes of an intrinsically disordered protein, Prostate-Associated Gene 4 (PAGE4). PAGE4 is a transcriptional coactivator that potentiates the oncogene c-Jun. Two kinases, namely HIPK1 and CLK2, phosphorylate PAGE4 generating variants phosphorylated at different serine/threonine residues (HIPK1-PAGE4 and CLK2-PAGE4, respectively) with opposing functions. While HIPK1-PAGE4 predominantly phosphorylates Thr51 and potentiates c-Jun, CLK2-PAGE4 hyper-phosphorylates PAGE4 and attenuates transactivation. To understand the underlying mechanisms of conformational diversity among different phosphoforms, we have analyzed their atomistic trajectories simulated using AWSEM forcefield and the energy landscapes were elucidated using ELViM. This method allows us to identify and compare the population distributions of different conformational ensembles of PAGE4 phosphoforms using the same effective phase space. The results reveal a predominant conformational ensemble with an extended C-terminal segment of WT PAGE4, which exposes a functional residue Thr51, implying its potential of undertaking a fly-casting mechanism while binding to its cognate partner. In contrast, for HIPK1-PAGE4, a compact conformational ensemble enhances its population sequestering phosphorylated-Thr51. This clearly explains the experimentally observed weaker affinity of HIPK1-PAGE4 for c-Jun. ELViM appears as a powerful tool especially to analyze the highly-frustrated energy landscape representation of IDPs where appropriate reaction coordinates are hard to apprehend.

## Introduction

Contrary to Anfinsen’s hypothesis that structure defines protein function,^1^ it is now increasingly evident that a significant fraction of the human proteome is composed of intrinsically disordered proteins (IDPs) that lack rigid 3D structure.^2–5^ Despite the lack of well-defined structure under physiological conditions at least in vitro, IDPs are involved in a large number of biological functions ranging from gene-regulation, molecular recognition, signal transduction and intracellular information processing events^6–8^ underscoring their functional significance.

IDPs can adopt a huge repertoire of conformations, because of their high backbone flexibility and inherent plasticity,^9–13^ and hence, exist as conformational ensembles. Thus, the conformational search process of an IDP involves a higher-dimensional phase space than that of a structured/ordered protein. While enormous research efforts have been spurred to understand the classical “minimally-frustrated” or “funnel-like” energy landscapes for structured proteins,^14–16^ understanding “highly-frustrated” or “weakly-funneled” nature of IDPs has remained far more challenging due to their large-scale conformational changes. ^17–20^ Furthermore, compared to ordered proteins, IDPs are relatively more susceptible to post-transitional modifications, especially phosphorylation,^21–23^ which has been shown to impinge on their conformational dynamics.^7,24–26^ In several cases, such conformational switches induced by site-specific phosphorylation has been implicated in disease pathology.

Concomitant with advances in experimental techniques typically used to investigate IDPs, such as nuclear magnetic resonance (NMR), small-angle X-ray scattering (SAXS), and single-molecule fluorescence resonance energy transfer (smFRET),^27–31^ over the past two decades, a number of computational techniques have been developed, causing an explosion in research on the IDPs.^32,33^ Nonetheless, capturing the detailed conformational dynamics of IDPs at an atomistic length scale still remains a challenge both experimentally and computationally. More specifically, the challenges are two-fold: (i) Generating high-resolution ensemble of highly flexible IDPS is computationally expensive in most cases as a large number of conformational changes are associated with an IDP; (ii) Due to the numerous degrees of freedom in complex IDPs, mechanistic interpretation of trajectories from computer simulations is a tough call, if not impossible. The former challenge imparts the sampling issues concerning the huge phase-space of IDPs. Canonical atomistic MD simulations are often found futile in simulating IDPs due to inadequate statistical sampling. Ideally, sampling an IDP requires long-time simulations, and those are computationally highly expensive. To address the sampling issues, different coarse-grained (CG) models,^34,35^ trained molecular simulations by experimental inputs,^36–38^ *de-novo* enhanced molecular dynamics simulations techniques are developed and found quite useful in rapid sampling of the IDP phase-space.^39–41^ While most of computational techniques focus on the sampling issues of IDPs, which of course is essential, mechanistic interpretation of trajectories from computer simulations still remains a major challenge.

Among IDPs that have been extensively studied, Prostate-associated gene 4 (PAGE4) is one of the most well-characterized IDPs in terms of its biology and its conformational preferences and dynamics.^25,42,43^ PAGE4 is a stress-response protein that acts as a transcriptional coactivator. In prostate cancer (PCa) cells, PAGE4 potentiates transactivation of c-Jun that heterodimerizes with c-Fos to form the Activator Protein-1 (AP-1) complex.^44^The stress-response kinase, homeodomain- interacting protein kinase 1 (HIPK1), phosphorylates PAGE4 predominantly at Thr51 with a minor fraction of the ensemble being phosphorylated at Ser9.^45^ On the other hand, CDC-like kinase 2 (CLK2) hyper-phosphorylates PAGE4 (CLK2-PAGE4), and in contrast to HIPK1-PAGE4, CLK2-PAGE4 inhibits c-Jun activity.^43^ Consistent with the biochemical data, NMR studies indicated that HIPK1-PAGE4 exhibits stronger binding to AP-1 than that of CLK2-PAGE4. Furthermore, molecular dynamics simulations using Atomistic Associative Memory, Water Mediated, Structure and Energy Model (AAWSEM) revealed that this difference in binding affinity may be attributed to the phosphorylation-induced conformational changes adopted by HIPK1-PAGE4 and CLK2-PAGE4. Confirming the results from molecular simulation, SAXS, smFRET results show that HIPK1-PAGE4 exhibits a relatively compact conformational ensemble, whereas the conformational ensemble of CLK2-PAGE4 appears more expanded.^43^

Given the complexity and 3D spread of the conformational space of an IDP as highly disordered as PAGE4,^27,46^ deriving an ensemble average picture representing its conformational plasticity may represent a rather crude approach. By this approach, a functional conformational ensemble might stay underestimated. Therefore, visualization of the entire conformational ensemble map is important that can help identify the mechanistic interplay between different conformational clusters and their functional implications. With this goal in mind, here, we have adopted ELViM approach, a unique and useful tool to discern the rough energy landscapes of IDPs and rich characteristics of IDP dynamics.^47^ Given a data set of conformations and using an appropriate metric, ELViM calculates a matrix of internal distances between all pairs of conformations. This matrix, which represents the data set in the high dimensional phase space, is then projected into an effective 2D phase space. The analysis of this final energy landscape representation allows us to infer and establish a connection with functional mechanisms.

## Results

The simulation trajectories of PAGE4 consists of 10^4^ configurations of each of the three variants: Wild Type (WT), CLK2, and HIPK1, which were described in a previous study.^42^ The sequence of PAGE4 and its phosphorylated and non-phosphorylated (WT) residues are indicated in Figure 1.a. In this work, we have employed our newly developed ELViM approach for the first time to not only visualize and understand the higher dimensional conformational phase-space of a highly flexible IDP, but also to compare the different conformational ensembles. ELViM being a reaction coordinate-free method nicely provides an unprecedented and intuitive 2D visualization of the multidimensional phase-space. The effective 2D phase-space of all PAGE4 conformers is given by an approximately circular disk, shown in Figure 1.b, in which, each structure is represented by a point. One can see the conformational states of WT-PAGE4 and HIPK1-PAGE4 occupying similar regions of the phase space, while CLK2-PAGE4 occupies a region in the phase-space that is distinct from that occupied by the WT and HIPK1 ensembles. However, ELViM brings all conformational ensembles of each conformer under the same effective phase-space region where the ensemble overlaps and distinctions can be discerned simultaneously.

**Figure 1:**
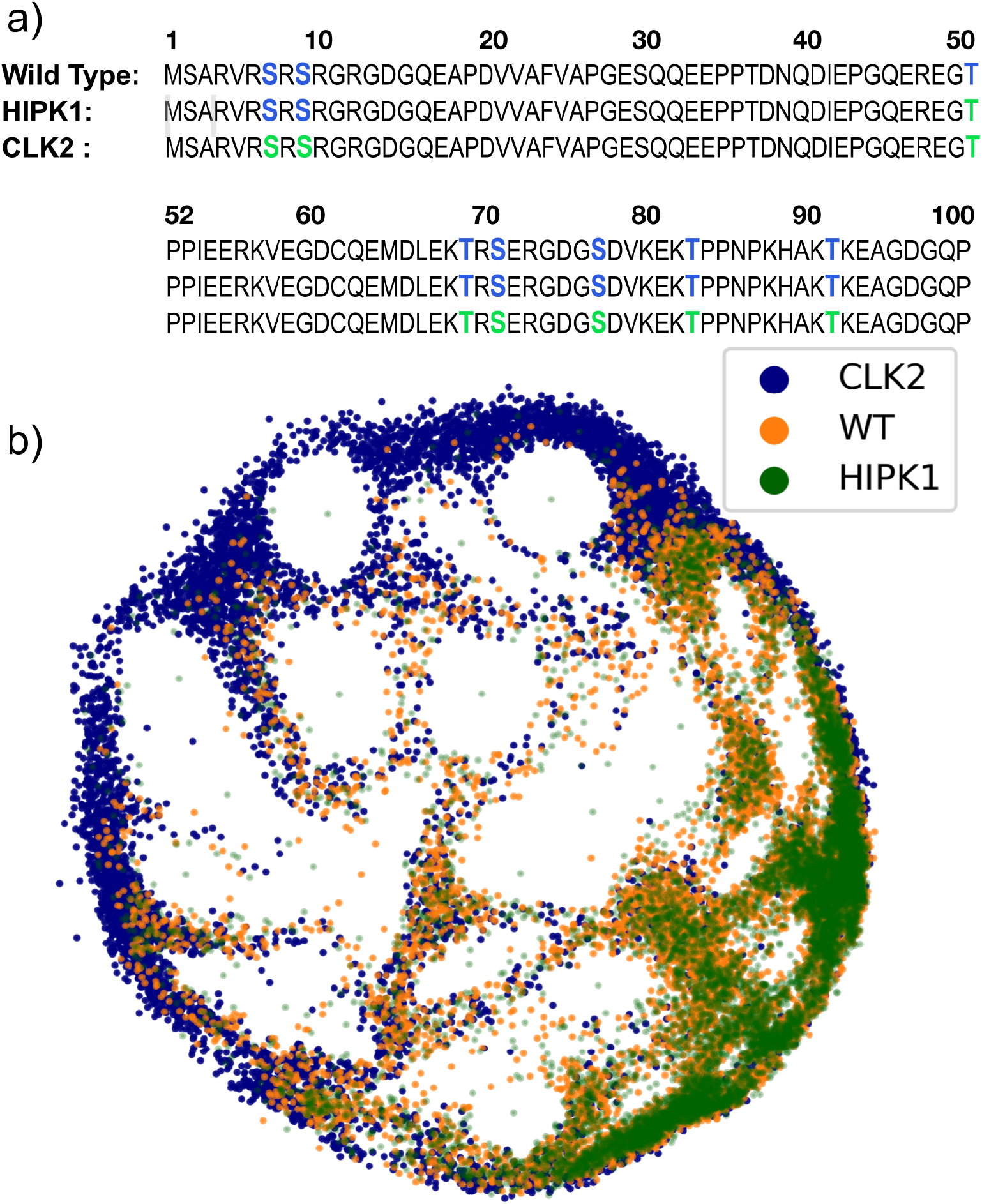
PAGE4 Sequences and 2D ELViM Projection. (a) PAGE4 Wild type, HIPK1 and CLK2 sequences, in which the non-phosphorylated and phosphorylated residues are shown in blue and green respectively. (b) Each configuration of each type of phosphoform is represented as a point in the ELViM’s effective 2D representation.

### Validation of 2D phase space

If the studied system is not very complex, ELViM can provide a meaningful energy landscape of the structures. One way to validate ELViM representation is to probe measurable variables and determine if they are well behaved, *i.e.*, if they vary continuously. Figure 2 shows the radius of gyration (*R_g_*) of different regions of the protein for each state where different conformations are enveloped. This analysis is consistent with our early simulation measurements and indeed captures the dynamical changes in the PAGE4 ensemble. By separately accounting for N- and C-terminal regions, we previously observed that both N- and C-terminal regions have a loop forming tendency.^43^ However, the present study revealed that the N-terminal part more frequently adopts a compact turn-like structure compared to the C-terminal region, Figure 2(b) and (c). This clearly implies that the N-terminal half dynamically contributes more to the overall compaction of PAGE4. Such N- and C-terminal conformational dynamics of PAGE4 and its different phophoforms were also monitored in an early smFRET experiments where the N terminal size-variability was measured by probing the distance between residues 18 and 63.^25^ Interactions between residues 18 and 63 and between residues 63 and 102 were calculated for every conformation in the ELViM projection and the results show good agreement with experimental observations while validating the well behaved representation of the data (see Fig.SI.1). Consistent with the present ELViM analysis, our previous observations employing smFRET also indicated that the C-terminal region of PAGE4 is less dynamic in terms of its conformational preferences in response to phosphorylation compared to its N-terminal half. While the phase-space of *R_g_* characterizes the overall macroscopic size-variability of this IDP and its different phosphoforms, how the segmental loop forming tendency evolves microscopically in each ensemble is yet to be determined.

**Figure 2:**
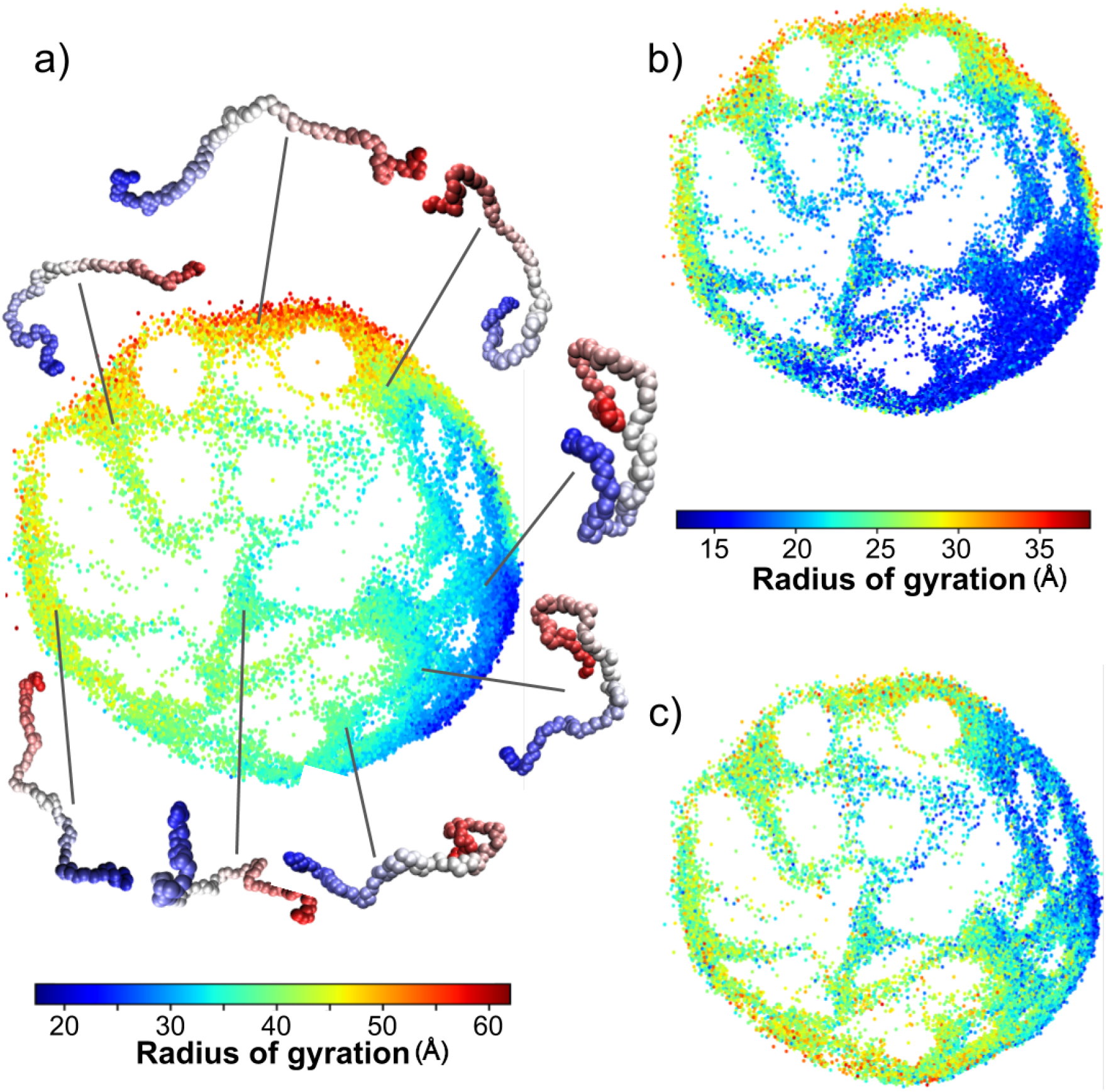
Structures as a function of radius of gyration. Each conformation is colored as function of radius of gyration of (a) the entire protein, (b) of the N-terminal part (residues 1-51), and (c) of the C-terminal part (residues 52-102). The shown structures are typical examples of each area of the 2D effective phase space, in which N- and C-terminal are shown in red and blue, respectively.

### Analysis of each PAGE4 phosphoform

Once all conformations of the data set are projected into the 2D effective phase-space, one can start analyzing each ensemble data separately. Because all conformations are described under the same overall reference frame, comparisons between their behavior can be carried out simultaneously. A straightforward analysis is to obtain from the 2D density of states of each pixel *i* for each PAGE4 conformer 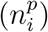, which is shown in Figure 3. The density of states (DOS) demonstrates that WT- and HIPK1-PAGE4 populate very distinct areas of conformations compared to CLK2-PAGE4. Also, CLK2-PAGE4 being a more expanded variant, its disorderedness and diminished affinity for c-Jun were very well-characterized in previous biophysical experimental measurements and simulation studies.^25,27,42,43,45^ Furthermore, while WT- and HIPK1-PAGE4 can bind to c-Jun, CLK2-PAGE4 has significantly lower binding affinity for c-Jun.^25,45^Thus, in the current study, we were more interested in analyzing and comparing the conformational heterogeneity between WT- and HIPK1-PAGE4. Interestingly, all previous measurements employing smFRET, NMR, and small-angle X-ray scattering (SAXS) showed that HIPK1-PAGE4 is slightly more compact than WT-PAGE4^25^ Subsequently, experimental measurements also found that in comparison to WT, HIPK1-PAGE4 has reduced binding affinity for c-Jun, which affects its transactivation of target genes.^45^ Therefore, to understand the differences between WT- and HIPK1-PAGE4, we compared their DOS maps where the effective phasespace appears very similar and in both the maps, we identify three distinct conformational clusters designated as: A, B, C in Figure 3.a and 3.b.

**Figure 3:**
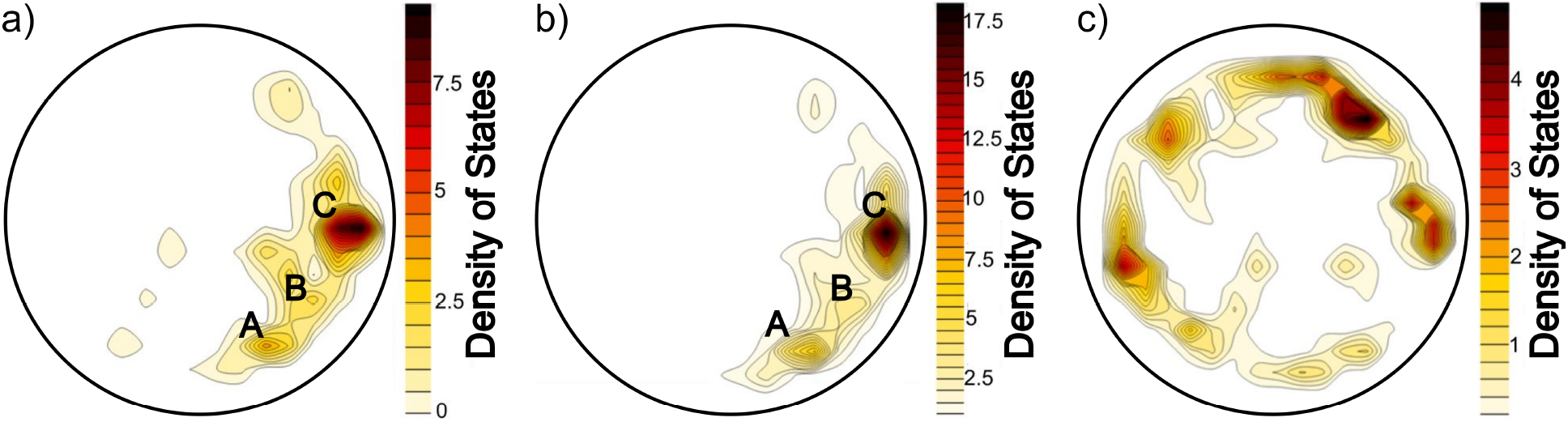
Density of states for each phosphoform. Each PAGE4 data set can be analyzed individually using the same 2D effective phase space shown in Figure 1.b., from which the density of sates is calculated. (a) WT-PAGE4, (b) HIPK1-PAGE4, and (c) CLK2-PAGE4.

A detailed analysis of these regions can provide insight into the overall features of these states and help in explaining the microscopic origin of the increased compaction in HIPK1-PAGE4 than that seen in WT-PAGE4 as determined by SAXS measurement.^25^ Region A is the highly populated stable region for WT- and HIPK1-PAGE4 in the entire phase-space. It isolates the most compact conformations where the compaction of both the N-terminal (1-51) and C-terminal (52-102) result in lowering of overall (*R_g_*) value as shown in Figure 2. Regions B and C also have a compact N-terminal region but they possess a more labile and extended C-terminal fraction. The contact maps associated with these regions corroborate this compactness feature. For a given conformation, a contact between residues *i* and *j* is considered formed if *r_ij_* ≤ *r_c_*, where *r_c_* is a cut-off contact distance. In this study *r_c_* = 8 Å. For a given set of structures, a contact is considered as formed if it occurs in at least 50% of the states. Figure 4 shows the contact map of WT-PAGE4 and HIPK1-PAGE4 in regions A, B and C defined in Figure 3.a and b. They show contacts between residues involved in the N-terminal portion, characteristic of loop formation. As the system goes from region A to B and C, one can identify a progressive C-terminal region extension. Typical structures of these regions are shown in Figure 2 and 5.a, 5.b, and 5.c.

**Figure 4:**
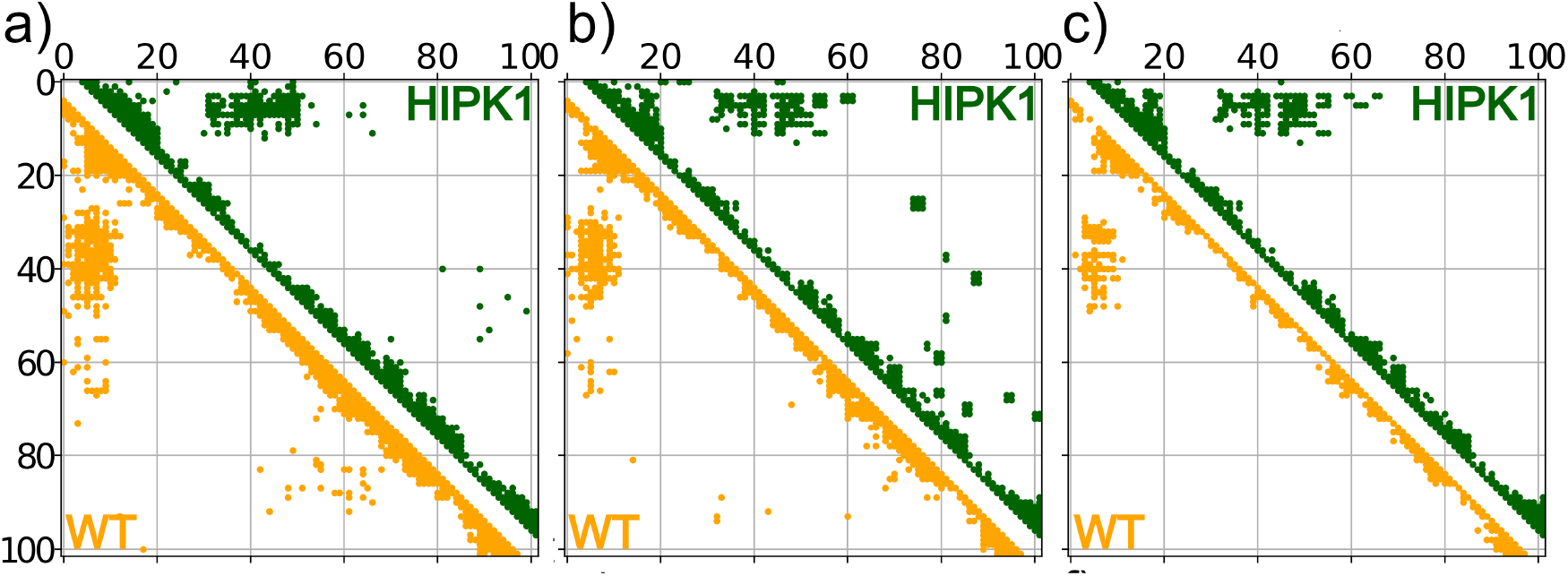
Contact maps for WT-PAGE4 and HIPK1-PAGE4. A residues *i* and *j* are defined in contact if they are within a distance *r_c_* (*r_ij_ ≤ r_c_*, with *r_c_* = 8. Å). WT-PAGE4 and HIPK1-PAGE4 structures were selected from regions, A, B, and C shown in Figure 3. A contact between residues *i* and *j* is considered formed if it occurs in at least 50% of the selected states. The contact maps for regions A, B, and C, are shown in (a), (b), and (c), respectively. In (c), there is no loop-forming contacts in region C-terminal segments, which indicates that this segment is in its extended form.

**Figure 5:**
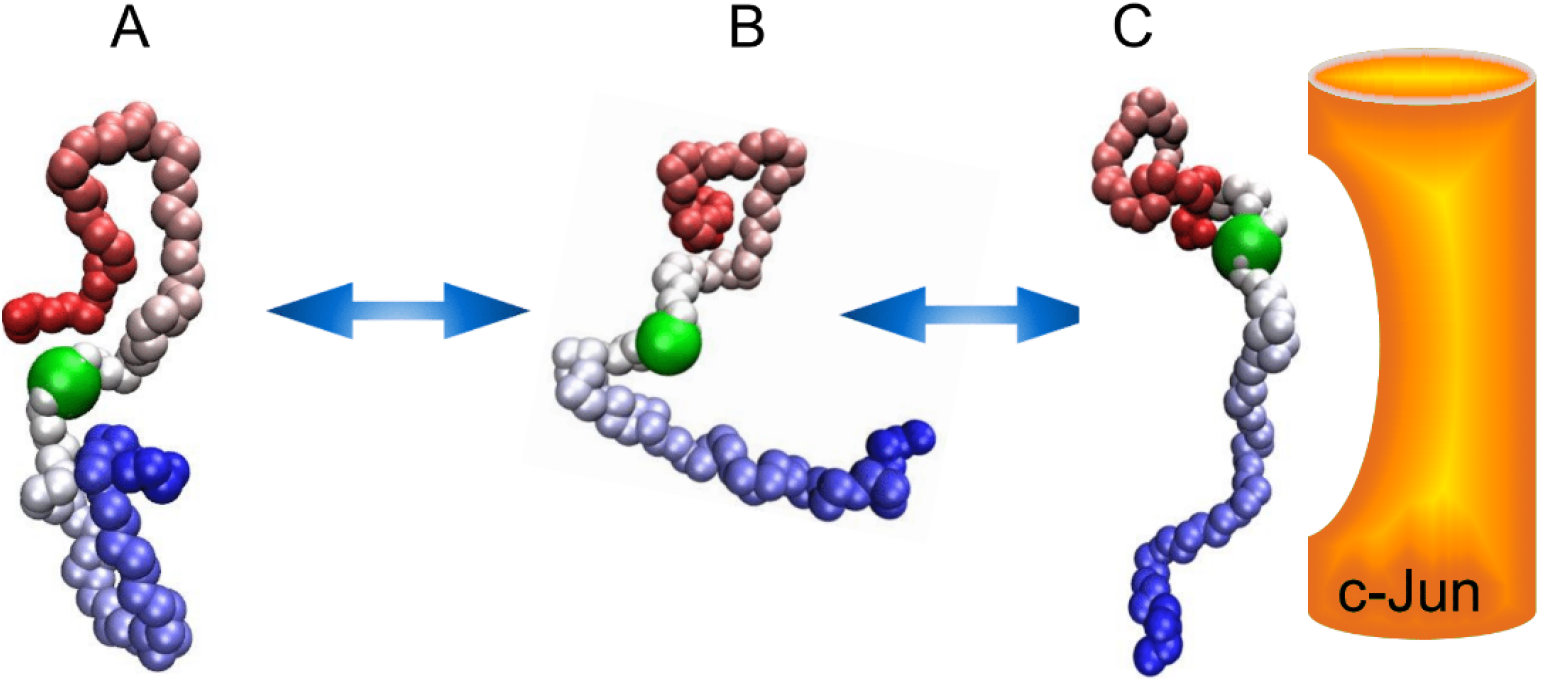
Wild Type PAGE4 Mechanism. Suggested fly-casting mechanism of WT-PAGE4, in which the C-terminal portion undergoes a conformational extension, from states A to B and C, allowing it to doc and activate the c-Jun complex. The green highlighted residue corresponds to Thr-51.

### The Fly-casting of Wild Type PAGE4

Our contact map analysis correlates well with earlier NMR spectroscopy results, which show that PAGE4 has local and long-range conformational preferences that are significantly perturbed by the site-specific phosphorylation at Thr51.^27^ It is evident that in WT PAGE4, these preferential interactions are clearly manifested in the N-terminal loop formation, keeping C-terminal relatively extended 3, and exposing the central acidic region where c-Jun binds.^27^ Our results suggest that such favorable WT-PAGE4 conformations facilitating c-Jun docking to occur are associated with states observed in the region C in Figure 3.a. The high stability of region A, the structural similarity between the conformations of A and C regions, together with intermediates from B region, offer a persuasive argument for a functional mechanism. As we move beyond the stable basin of region A, the C-terminal segment progressively becomes extended. It populates region B, finally reaching region C, which allows better exposure of the central acidic region and its close interaction with c-Jun complex, as depicted in Figure 5. This population-shift towards region C is likely to ensure PAGE4 functioning by extending its higher capture radius, exposing the binding site to c-Jun, but still retaining its moderate flexibility by a partial loop formation. The observation of partial loop formation hints at a fly-casting motion that is frequently observed in the conformational dynamics of disordered proteins.^48,49^

### The Occlusion of HIPK1-PAGE4

Since the 2D conformational phase-space of WT- and HIPK1-PAGE4 are very similar, it would be expected that they would have the same binding mechanism to c-Jun. However, experimental results indicate that HIPK1-PAGE4 has a lower binding affinity for c-Jun.^27^ One possible reason for the lowered affinity of HIPK1--PAGE4 for c-Jun *in vitro* may be due to the more compact nature of the conformational ensemble of HIPK1 than WT-PAGE4. Here, ELViM plays a very important role by capturing the population shift towards region A where both N- and C-terminal regions contribute to form a compact core sequestering the central acidic regions. Such a population shift in HIPK1-PAGE4 renders an occlusion effect barring HIPK1-PAGE4 from interacting with c-Jun, which may reduce their binding affinity as found in experiments.^45^ To qualitatively monitor this population shift mechanism one can estimate the free energy from ELViM 2D representation as 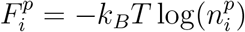 using the DOS shown in Figure 3. Probing the points along the path that connects the basins A, B, and C, the approximate free energy differences can be inferred, 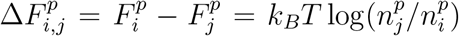. The approximate free energy profile for WT-PAGE4 and HIPK1-PAGE4 are shown in Figure 6. HIPK1-PAGE4 presents higher energy barriers and higher free energy difference between basin A and C when compared with WT-PAGE4. Based on both kinetic and thermodynamic aspects, it makes functional conformations in region C less likely to be probed in HIPK1-PAGE4. The energy differences are in the order of *K_B_T* or less and hence, it may be argued that these are too small to make a difference in the mechanisms. However, it should be noted that the simulation model is coarse-grained so the values for the energy barriers and free energy difference are qualitative.

**Figure 6:**
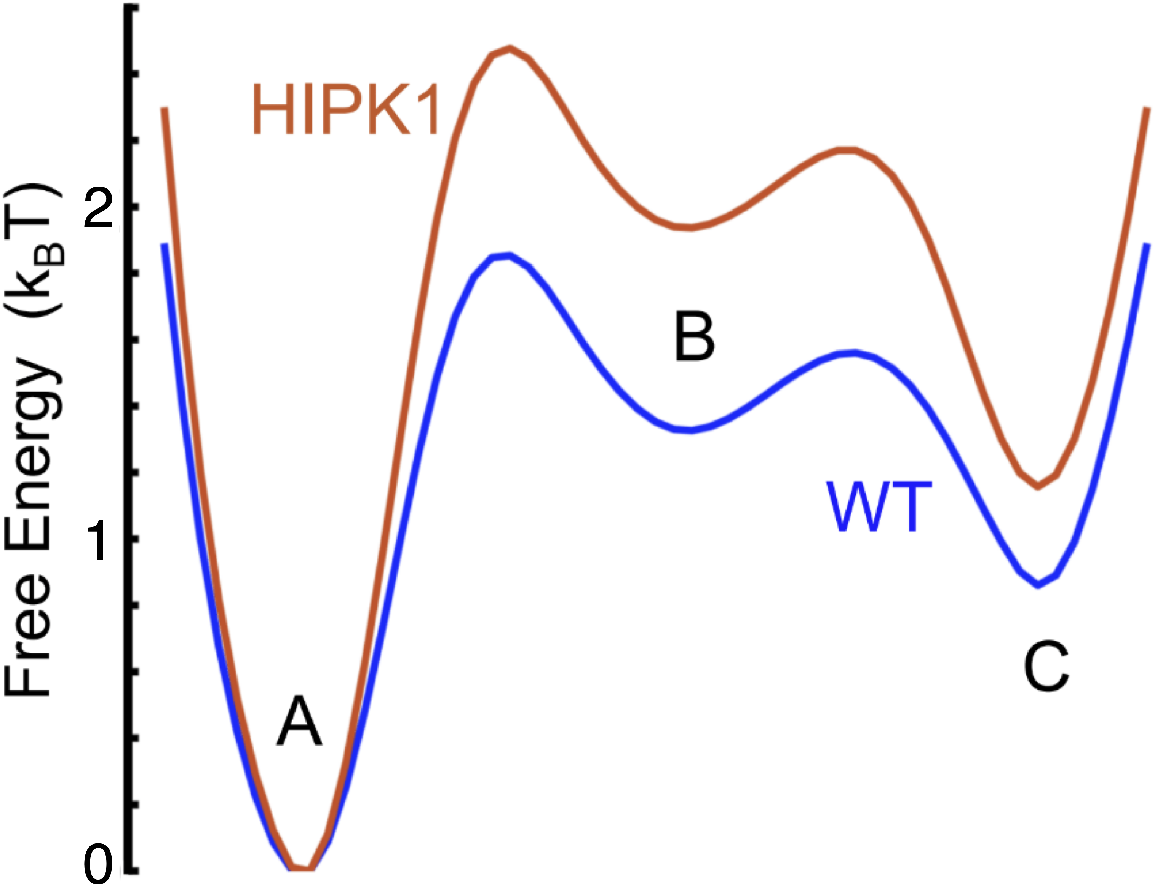
WT- and HIPK1-PAGE4 Free Energies. Comparison of the free energy profiles based on the density of states along the path from regions A to B, and C. HIPK1-PAGE4 profile presents a higher free difference between A a C ensembles and a higher transition from A to C when compared with WT-PAGE4.

### Central Role of Thr-51 in Maneuvering Conformational Equilibrium

Biological macromolecules, including IDPs, adopt a diverse ensemble of conformations for which distinct functions have evolved. The populations of conformational ensembles preserve a dynamic equilibrium as we currently find among A, B, and C population regimes. It is clear from our structural analysis that the underlying differences between WT- and HIPK1-PAGE4 have emerged because of the flexibility of the C-terminal motif. The conformational flexibility of the C-terminal motif again depends on its preferential interaction with the acidic region centered on Thr-51. Thus, Thr-51 serves as a controlling anchor, which has the potential to assemble both N- and C-terminals making 8-ring like structures (region A). In HIPK1-PAGE4, the population-shift occurs from region C to A which is highly enriched by such 8-ring like structures where Thr-51 is highly buried by the neighboring residues. This feature is readily inferred by measuring the radius of gyration (*R_g_*) of segments around Thr-51. We calculate *R_g_* in segments from 51 − *N* to 51 + *N* for all the conformations in the ELViM 2D representation. This result is also consistent with recent NMR spectroscopic measurements. These measurements revealed that the largest chemical shift occurs at Thr-51, suggesting that Thr-51 envelopes a number of long-range neighbors in the central region. Thus, it becomes more compact and more negatively charged in the HIPK1-PAGE4 version.

Figure 7 shows *R_g_* in the interval [26,76] (*N* = 25). The data indicates that the region in which HIPK1-PAGE4 has the highest density of states (region A), corresponds to the lowest *R_g_*, *i.e.* the segment containing Thr-51 in its center is expected to be strongly packed which is in good agreement with experimental results. Figure SI.2 shows *R_g_* in the intervals [41, 51], [31, 71], and [16, 86] (*N* = 10, 20, and 35, respectively), which indicates similar behavior as in Figure 7.

**Figure 7:**
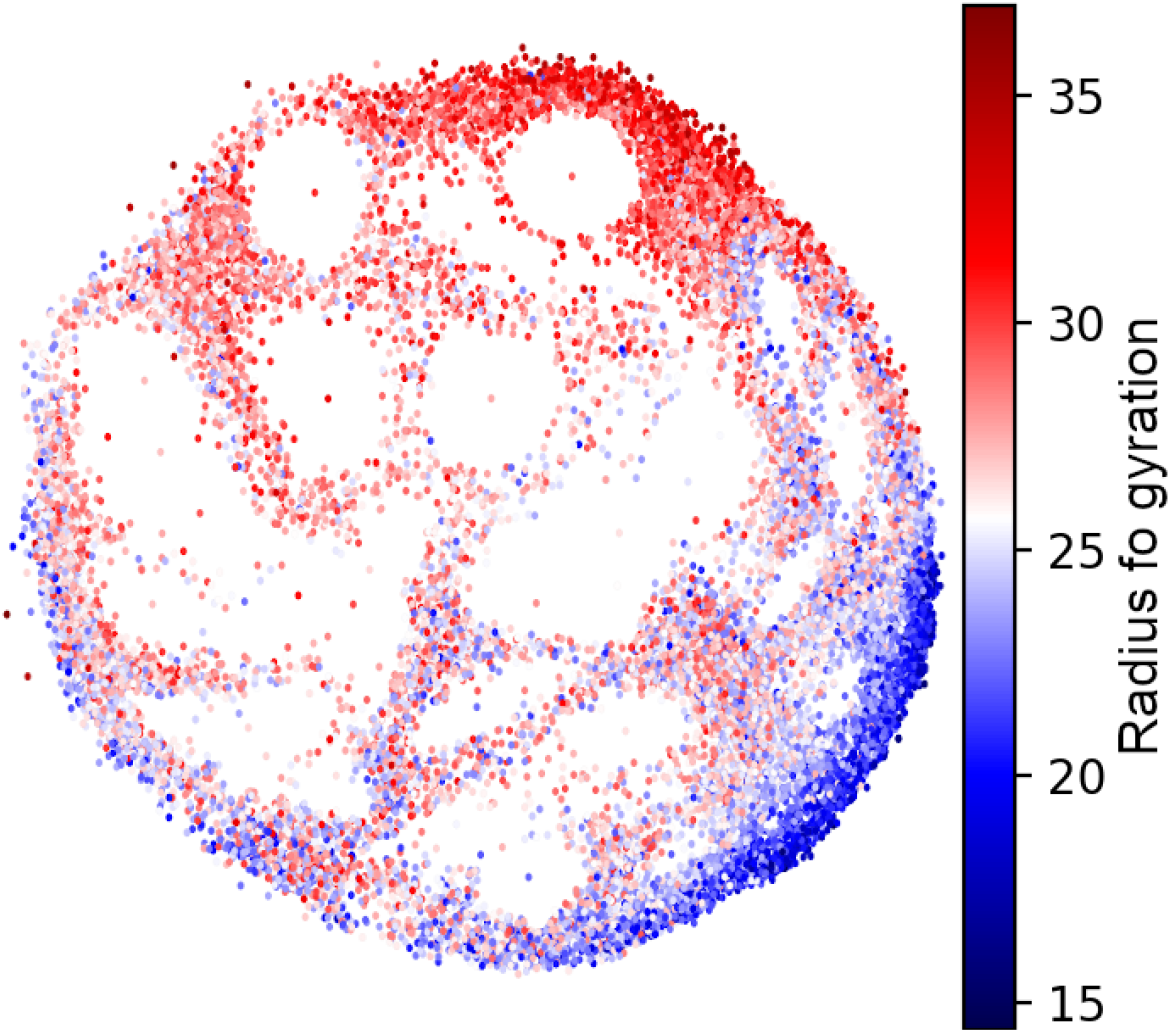
Role of Thr51. *R_g_* of the segment in the interval [51-*N*, 51+*N*] with *N* = 25. In region A defined in Figure 3.b, *R_g_* is the lowest, which suggests that Thr51 is buried the most, in agreement with experimental results.^25^

**Figure 8:**
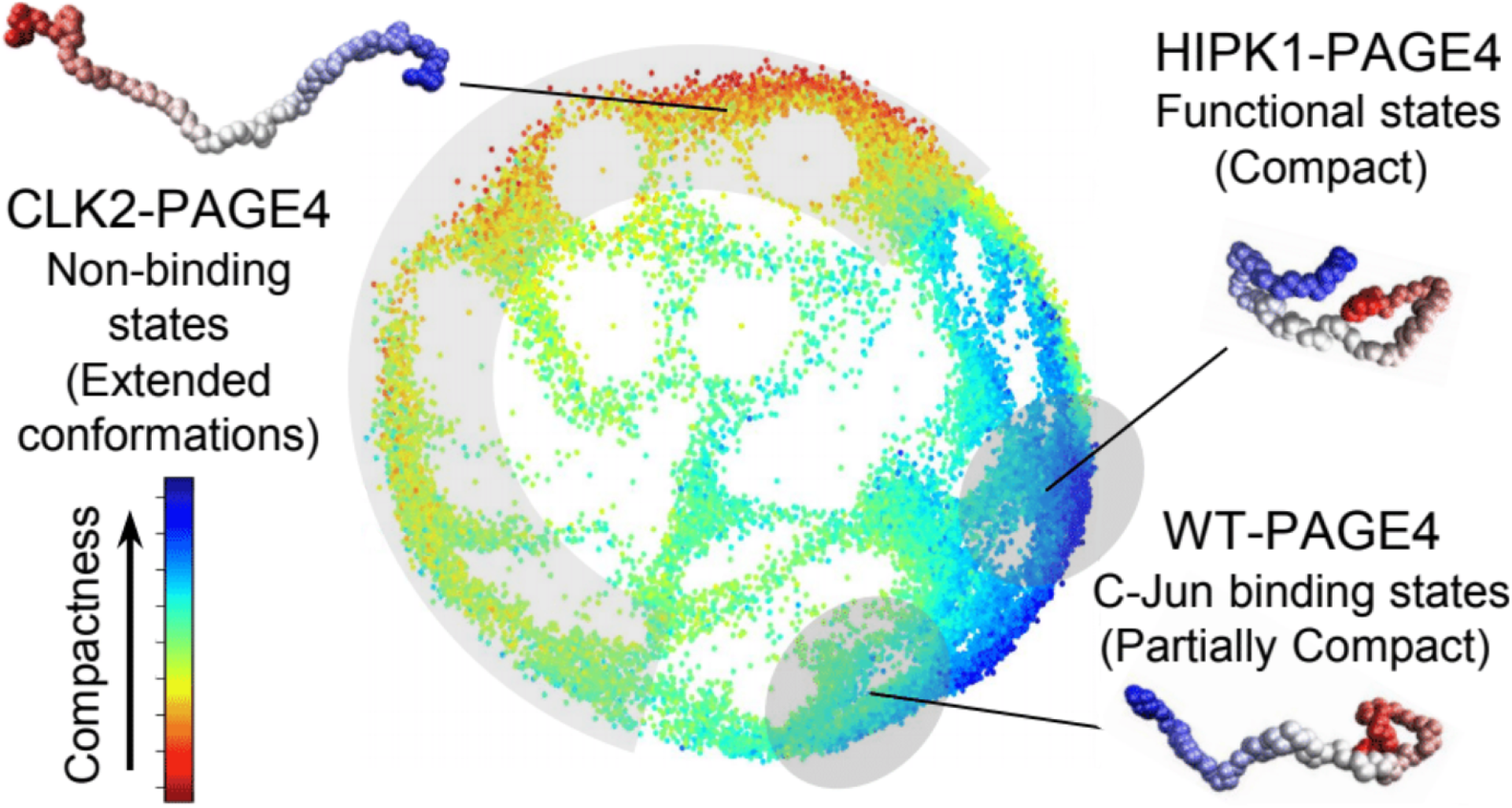
For Table of Contents Only.

## Conclusion

Energy landscapes of IDPs are typically hard to investigate due to the lack of a single native reference conformation. Moreover, due to their plasticity, IDPs tend to explore an unusually larger conformational phase-space when compared with classical minimally-frustrated proteins, and hence, putative reaction coordinates are practically non-existent. Even when *a posteriori* reaction coordinates are found, they have the potential to mask the richness of the dynamics. In contrast, ELViM represents complex multidimensional landscapes in a simplified two-dimensional phase-space using a purely data-driven approach, in which no assumption of reaction coordinates or a reference conformation is needed. This reaction-free approach is particularly useful in the case of IDPs. The technique also allows comparisons between proteins studied under different conditions in a single framework.

In the present manuscript, the method was applied to a highly intrinsically disordered protein, PAGE4. The ELViM representation of the data is in excellent agreement with the experimental results obtained using a variety of biophysical techniques. More specifically, the previous computational results, such as the likelihood of loop formation due to phosphorylation and the structural characteristics of the different PAGE4 conformational ensembles, highlight the virtues of employing ELViM to gain deeper insight. In particular, new details of the mechanisms involved in PAGE4 variants are unveiled applying ELViM. For example, hitherto forth, the WT fly-casting mechanism was thought to be a plausible binding mechanism;^43^ however, the ELViM results suggest this may not be the case with HIPK1-PAGE4. The burial of Thr-51, which plays a key role in c-Jun binding, has also been elucidated by analyzing *R_g_* in the dominant structural basin of the WT- and HIPK1-PAGE4. Considering the ELViM approach to investigate PAGE4 as a paradigm, it is expected to work equally well for other IDPs, and may represent an invaluable tool to address such challenging systems.

## Methods

### Simulation data of PAGE4 from AAWSEM model

The simulation trajectories for all PAGE4 phosphoforms were obtained using the AAWSEM (Atomistic Associative memory, Water mediated, Structure and Energy Model) force field^50,51^. AAWSEM model is a multi-scale model combining full atomistic simulations of the local segments of proteins with a coarse-grained simulation of the full protein structures.^52^ The model has been shown to successfully fold proteins composed mainly of *α* helices^50^ and of *α*/*β* structures.^51^ Since the original AAWSEM potential leads to an overly compact PAGE4 ensemble, the non-bonded contact potential was modified for better reflecting the size distribution of a disordered protein. Details about the choice of parameters, as well as the simulated trajectories of PAGE4 was discussed in our previous paper.^42^Briefly, a total of three forms of PAGE4, WT-PAGE4, HIPK1-PAGE4, and CLK2-PAGE4, were segmented based on their experimentally determined functional motif annotated in the literature.^27^ Those segments were simulated with the CHARMM36m force field,^53^ which has a balanced treatment of ordered and disordered proteins. The clustered structures from the all-atom simulations were passed into the “memory terms” of the coarse-grained AWSEM potential. The entire proteins of PAGE4 were simulated under the constant temperature of 300K using the coarse-grained AWSEM potential. The electrostatic interactions were treated using the Debye-Hückel approximation.^54^ The electrostatic interactions were also scaled by 4.0 to achieve a relatively balanced strength compared to the other terms of the coarse-grained AWSEM potential. The phosphorylated residues of PAGE4 were modeled separately in the all-atom and coarse-grained simulations. HIPK1-PAGE4 was phosphorylated at two sites, Ser9 and Thr51. CLK2-PAGE4 was phosphorylated at eight sites, Ser7, Ser9, Thr51, Thr71, Ser73, Ser79, Thr85, Thr94.^25^ The phosphorylated residues were simulated by adding phosphoryl group onto the original residues in the all-atom simulation. In the coarse-grained simulations, those residues were simulated by replacing the phosphorylated residues with “super-charged” glutamic acid (Glu) residues.^55,56^ Our simulation quantitatively reproduced the size distribution of three phosphoforms of PAGE4 as measured by SAXS and smFRET experiments,^25^ as well as the change of local secondary structures upon phosphorylation, indicated by the NMR experiments.^27^

### Energy Landscape Visualization Method (ELViM)

The visualization of biological macromolecule landscapes is achieved by optimizing a 2D representation of distances between structures of a given data set of conformations.^47,57^ The method is based primarily on computing a matrix with effective distances between all pairs of structures. Such measurement is based on internal distances between every amino-acid in the protein. The method seeks to represent every state of the data set by a point in a 2D phase space, which aims to describe the computed effective distances in an optimal manner. The method goes beyond the usual one-dimensional representation, and it does not require a reference-conformation or a reaction coordinate. This approach relies only on structural information. Nevertheless, the states’ energies are implicitly accounted for, since they are strongly correlated with their molecular conformations.

The visualization method is based on four steps: (i) an ensemble of structures which, in general, are obtained through simulations is generated; (ii) the dissimilarity matrix is calculated by applying a metric throughout the simulated trajectory; (iii) a data processing procedure is carried out to cluster very similar structures into a single conformation; (iv) a multidimensional projection is performed and, then, the dissimilarity matrix is transformed to a two-dimensional (2D) projection. In summary, ELViM can automatically detect pathways and competing transition state ensembles without the need for an *a priori* reaction coordinate. Instead of having to intuitively design coordinates, which can involve visual inspection of large numbers of transition events, ELViM provides a means for efficiently mapping structurally distinct pathways during elaborate conformational rearrangement in multi domain, or multi component, assemblies.

In this work, we propose a slightly different analysis; we carry out the projection for different systems simultaneously, using the same phase space. In this case, 10,000 structures of each simulation separately were selected (WT, CLK2, and HIPK1); from these data we performed the steps (ii) to (iv) described above in order to obtain as results the effective 2D projection. The Information about which simulation each structure belongs to is preserved and can be labeled.

## Supporting information

Supplementary Results

## Acknowledgement

This research was supported by the Center for Theoretical Biological Physics sponsored by the NSF (Grants PHY-2019745 and CHE-1614101) and by the Welch Foundation (Grant C-1792). ABOJ was funded by FAPESP (São Paulo Research Foundation and Higher Education Personnel) Grant 2016/01343-7 and is currently a Robert A. Welch Postdoctoral Fellow. JNO is a Cancer Prevention and Research Institute of Texas (CPRIT) Scholar in Cancer Research. VBPL was funded by FAPESP Grant 2019/22540-3 and 2018/18668-1. SR acknowledges support from Science and Engineering Research Board (Grant SRG/2020/001295) and Department of Biotechnology, Govt. of India (Grant BT/12/IYBA/2019/12).

## Supporting Information Available

The following files are available free of charge.

- Filename: PAGE4-ELViM-SI.pdf. It contains the Support Figures referred to in the main text. Figure SI.1 shows ELViM projection conformations colored according to the smFRET probed distances. Figure SI.2 shows ELViM projection conformations colored by *R_g_* associated with different segments around Thr51.

