## Supplementary Results for "Exploring Energy Landscapes of Intrinsically Disordered Proteins: Insights into Functional Mechanisms"

### Supplementary Information for: Exploring Energy Landscapes of Intrinsically Disordered Proteins: Insights into Functional Mechanisms

Antonio B. Oliveira Junior,[†] Xingcheng Lin,[‡] Prakash Kulkarni,[¶] José N. Onuchic,[†] Susmita Roy*,[§] and Vitor B.P. Leite*,[||]

†*Center for Theoretical Biological Physics, Rice University, Houston, TX, USA*

‡*Department of Chemistry, Massachusetts Institute of Technology, Cambridge, MA, USA*

¶*Department of Medical Oncology and Therapeutics Research, City of Hope National Medical Center, Duarte, CA 91010*

§*Department of Chemical Sciences, Indian Institute of Science Education and Research Kolkata, West Bengal 741246, India*

||*Departamento de Física, Instituto de Biociências, Letras e Ciências Exatas, Universidade Estadual Paulista (UNESP), São José do Rio Preto, SP, Brazil*

.

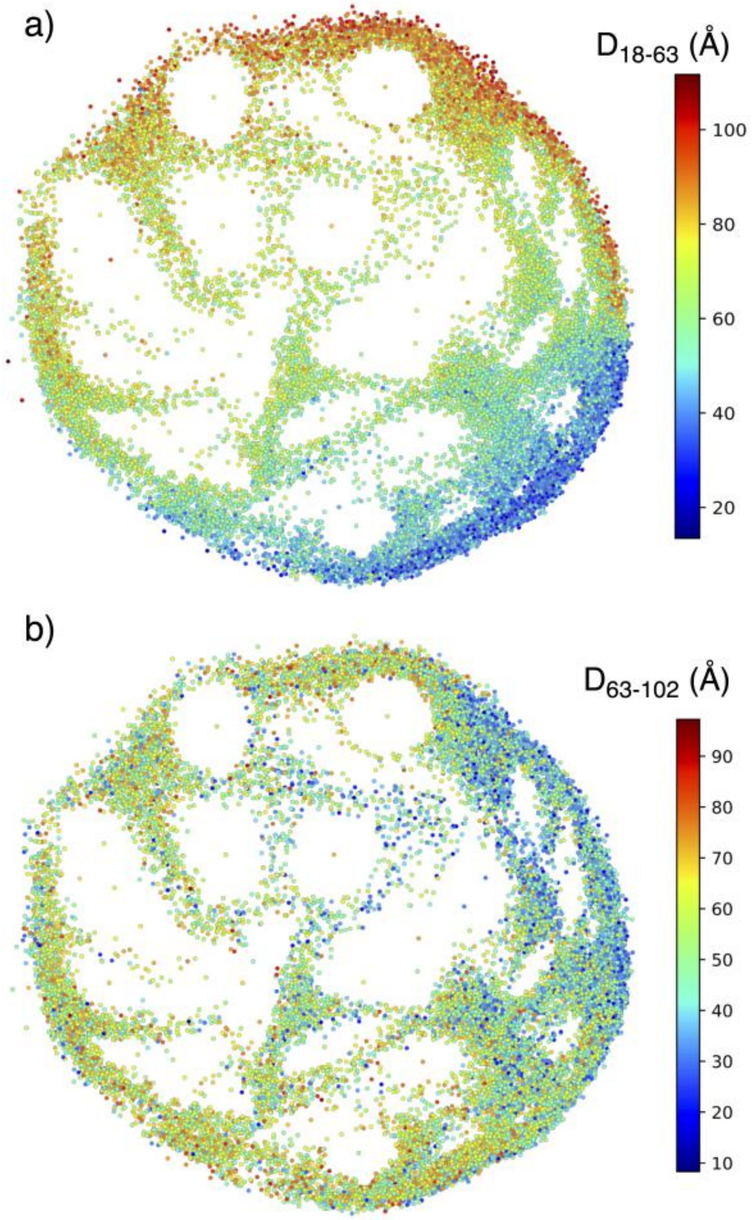


Fig. SI 1: The distances that measured by smFRET probes,[1] were calculated for each conformation in the ELViM projection, and colored accordingly. The distances were (a) between residues 18 and 63, and (b) between residues 63 and 102.

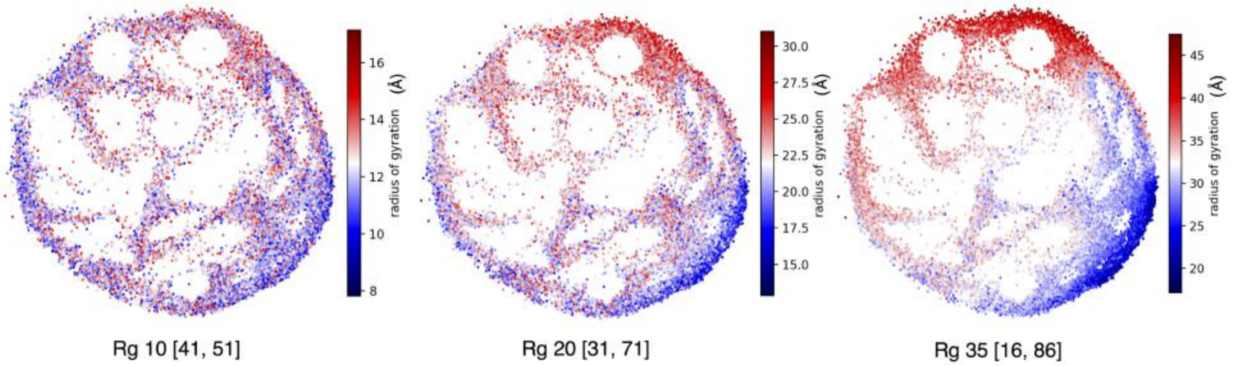


Fig. SI 2: ELViM projection conformations colored by $R_g$ associated with different segments around Thr51, which are in the interval [51-$N$, 51+$N$]. (a) $N = 10$, segment [41, 51], (b) $N = 20$, segment [31, 71], and (c) $N = 35$, segment [16, 86].
